# Farm yard manure enhances phosphate fertilizer use efficiency in different upland rice genotypes through mitigating aluminium toxicity

**DOI:** 10.1101/2020.05.13.088294

**Authors:** Pieterjan De Bauw, Erina Shimamura, Tovohery Rakotoson, Andry Andriamananjara, Mieke Verbeeck, Roel Merckx, Erik Smolders

## Abstract

Upland rice production on weathered soils is often constrained by phosphorus (P) deficiency and soil acidity. Farmyard manure application (FYM) can sharply enhance yields and agronomic P fertilizer (TSP) efficiency. We tested the hypothesis that rice genotypes differ in the extent of using organic P and offering distinct benefits under TSP-FYM combinations. Multiple field trials were conducted in the uplands of Madagascar, with factorial combinations of six genotypes, FYM, and TSP applications, with blanket N&K additions. Rice grain yields reached 6 t ha^-1^ after three years of TSP+FYM application, were lower when FYM or TSP were used separately, while crops failed under zero P input. Genotypic differences were inferior to the large treatment effects. Application of FYM increased soil pH and CaCl_2_-extractable P while decreasing CaCl_2_-extractable aluminium. An additional liming trial indicated that beneficial effects of FYM over TSP relate to liming effects. Genotypic ranking of yields and agronomic efficiency was inconsistent, without superior genotypes under FYM versus TSP. However, Chomrong Dhan and FOFIFA 172 showed superior yields under TSP+FYM. The FYM application lowers aluminium toxicity which overrules potential effects of organic P supply. Aluminium tolerance should be included when developing rice genotypes for low P tolerance in weathered soils.

## Introduction

Upland rice is often grown by subsistence farmers in Asia, Sub-Saharan Africa, and Central America, however, yield gaps of rice are larger in upland compared to lowland rice cropping systems (Chauhan, Jabran, & Mahajan, 2017). A large fraction of tropical upland systems are strongly nutrient depleted due to high soil weathering, persistent soil degradation and continuous cropping without external inputs (Stoorvogel et al. 1993). The weathered soils are dominated by sesquioxides who, combined with high soil acidity lead to low phosphorus (P) availability and low agronomic P use efficiency to the rice crop (Nishigaki et al., 2019). Therefore, upland rice production is often constrained by the combination of P limitations and soil acidity, logically aggravated by drought (Diagne et al., 2013; Mueller et al., 2012). For example, upland rice production in Madagascar contributes largely to the country’s total rice production, but a vast amount of the available upland area remains uncultivated as a result of these particular soil constraints (Andriamananjara et al., 2018; Raboin et al., 2014). To overcome P limitations in strongly weathered, acid upland soils, relatively large amounts of mineral P fertilizer are needed, which are often beyond reach to most common farmers (Nziguheba, 2007). Efforts are thus needed to increase the agronomic P fertilizer use efficiency. This can be achieved by soil management practices that increase the P availability to the crop, in combination with the use of low-P tolerant genotypes (Nziguheba et al., 2016).

The application of organic matter (OM) can enhance soil P availability. Several field trials with upland rice showed that farmyard manure (FYM) applications in the planting holes largely increase the agronomic P use efficiency in fields with added N&K fertilisers. (Andriamananjara et al., 2018, 2019). Organic anions released during OM decomposition compete with phosphate (PO_4_) sorption and improve availability of P from TSP. The FYM applications can temporarily enhance soil pH, thereby facilitating root proliferation and plant access to soil P while, at the same time, reducing PO_4_ sorption (Haynes and Mokolobate, 2001; Cong and Merckx, 2005). Finally, local high doses of FYM can enhance soil moisture content and reduce water stress. However, none of these factors have been experimentally disentangled to explain the beneficial effects on P uptake for upland rice.

Breeding for low P tolerant genotypes could be beneficial in upland rice cropping systems. The adaptation to P-limiting environments can derive either from an increased internal P utilization efficiency (PUE = [Tissue P concentration]^-1^)), i.e., how efficient the plant can use its tissue P to grow, or from an increased P acquisition efficiency (PAE = [Plant P uptake] * [Root biomass]^-1^), i.e. how efficient a plant can acquire P from the soil (Vandamme, Rose, Saito, Jeong, & Wissuwa, 2016). Genotypic variation exists in both PUE and PAE (Vandamme, Rose, et al., 2016; Vandamme, Wissuwa, et al., 2016; Wissuwa & Ae, 2001). Genotypic variation in PAE is related to a combination of differences in effective root surface area (Nestler & Wissuwa, 2016) per plant or in P acquisition strategies (Clark, 1983). For example, it is well established that plants can mobilize soil-adsorbed P by secreting root exudates, with different underlying P-solubilisation strategies from the inorganic P pool, or that plants can access organic P forms by the excretion of phosphatases (Richardson et al., 2011; Rose et al., 2013). The potential role of the organic P pool as a source of available P in tropical weathered soils has been emphasized in previous studies (Turner, 2006), however, information on the utilization of soil organic P by rice is scarce. One recent isotope dilution study with rice (Rakotoson, 2014), and an older study with ryegrass (Thibaud et al., 1988) and wheat (Osborne & Rengel, 2002) suggested that plant roots can locally absorb P from organic sources, likely by mineralization after phosphatase secretion or by interacting with microbial communities (Mehra et al., 2017). Potentially, rice genotypes may thus differ in the extent of using organic P and some genotypes may offer distinct benefits in FYM-TSP combinations compared to TSP only. However, only limited research has been conducted to assess the interaction between upland rice genotypes and the supply of organic soil amendments on the agronomic efficiency of P in highly weathered soils. The objectives of this study were to evaluate the interactions between rice genotypes and supply of FYM in P deficient soils and to identify soil chemical mechanisms that explain these interaction. We speculated that some genotypes may offer distinct benefits in FYM-TSP combinations. Multiple field trials were conducted in the uplands of Madagascar, with factorial combinations of six genotypes, FYM and TSP applications with blanket adequate N&K dosing and periodic irrigation, except for the first year. Specific attention was given to treatment effects on soluble P, Al, and soil pH by including an additional soil incubation trial and a mineral lime reference field trial for all genotypes.

## Materials and methods

### Experimental design

Field trials with factorial combinations of genotypes (Geno) and soil amendments were performed on three different, but adjacent fields with similar properties (Table 2). In field experiment 1, FYM-TSP-Geno interactions were tested for three subsequent seasons spread over three years, in field experiment 2, FYM-TSP-Geno interactions for two subsequent seasons and in field experiment 3, Lime-TSP-Geno interactions for one season. This combination of fields and seasons is referred to as an ensemble of six field trials, i.e. on three different fields, each over a period of, respectively, 3, 2, and 1 season. All plots were periodically irrigated unless in the first season of field 1. In addition, a laboratory incubation trial was set up to identify soil chemical processes explaining FYM-TSP interactions and to corroborate information from soil analysis of the field, sampled at the end of the field trials.

**Table 1.** Information of the six rice genotypes used in the field trials.

| Genotype Name | Genus | Varietal subgroup | Region of Origin | Varietal type | PAE<sup>a</sup> | RE<sup>b</sup> | References |
| --- | --- | --- | --- | --- | --- | --- | --- |
| NERICA 4 | *O. sativa* ×<br>*O. glaberrima* | *tropical japonica* | West Africa | Modern | low | low | De Bauw et al., 2019; Koide et al., 2013; Mori et al., 2016; Saito et al., 2015 |
| CG 14 | *O. glaberrima* | - | West Africa | Traditional | high | low | Koide *et al.* 2013<br>Wissuwa (personal communication) |
| DJ 123 | *O. sativa* | *aus* subspecies | Bangladesh | Traditional | high | high | Mori et al., 2016; Saito et al., 2015 |
| Chomrong Dhan | *O. sativa* | *temperate japonica* | Nepal | Traditional | NA | NA | Raboin *et al.* 2014 |
| FOFIFA 172 | *O. sativa* | *tropical japonica* | Madagascar | Modern | NA | NA | Raboin *et al.* 2014 |
| FOFIFA 173 | *O. sativa* | *tropical japonica* | Madagascar | Modern | NA | NA | Raboin *et al.* 2014 |

NA Data not available. <sup>a</sup>Phosphorus Acquisition Efficiency, expressed as total P uptake in mg at low P availability in the soil. <sup>b</sup>Root Efficiency, expressed as total P uptake per unit root size (either root biomass in g or root surface area in cm<sup>2</sup>).

**Table 2.** Information on soil properties of unamended (control) soil in the field and soil incubation experiments. Mean and standard deviations are presented when applicable (n=4 for Field 1 and 2; n=3 for the incubation experiment). Field experiment 3 was conducted at the edge of Field experiment 2, and hence soil properties are similar.

| Experiment | Field experiment<br>1 | Field experiment<br>2 | Field experiment<br>3 | Incubation<br>experiment<sup>§</sup> |
| --- | --- | --- | --- | --- |
| Start year | 2016 | 2017 | 2018 | 2019 |
| Duration | 3 seasons | 2 seasons | 1 season | 99 days |
| Coordinate | 19°12'59.3"S<br>47°27'10.6"E | 19°13'01.5"S<br>47°27'09.2"E | 19°13'00.4"S<br>47°27'09.5"E | 19°12'59.3"S<br>47°27'10.6"E |
| *Soil properties* |  |  |  |  |
| Soil classification<sup>a</sup> | Ferralsol | Ferralsol | Ferralsol | Ferralsol |
| Bulk Density (g cm<sup>3</sup>) | 0.98 | - | - | - |
| Soil texture<sup>b</sup> | Silty clay | Clay | ~ | NA |
| Sand (%) | 1.93±1.66 | 0.64±1.28 | ~ | NA |
| Silt (%) | 50.8±2.6 | 39.0±7.9 | ~ | NA |
| Clay (%) | 47.2±2.9 | 60.3±9.0 | ~ | NA |
| SOC<sup>c</sup> (%) | 3.0±0.5 | 3.0±0.3 | ~ | 3.3±0.1 |
| Soil pH<sub>CaCl2</sub><sup>d</sup> | 4.3±0.1 | 4.4±0.1 | ~ | 4.3±0.1 |
| eCEC<sup>e</sup> (cmol<sub>c</sub> kg<sup>-1</sup> soil) | 2.1±0.3 | 1.7±0.2 | ~ | 2.3±0.1 |
| P-total<sup>f</sup> (mg kg<sup>-1</sup> soil) | 680±30 | 550±10 | ~ | 620±10 |
| N-total<sup>c</sup> (mg kg<sup>-1</sup> soil) | 2029±360 | 1690±170 | ~ | 2220±90 |
| P-ox<sup>g</sup> (mg kg<sup>-1</sup> soil) | 26.7±4.1 | 17.3±1.9 | ~ | 29.3±3.3 |
| Fe-ox<sup>g</sup> (mg kg<sup>-1</sup> soil) | 1940±130 | 1620±80 | ~ | 1720±50 |
| Al-ox<sup>g</sup> (mg kg<sup>-1</sup> soil) | 2680±210 | 2650±90 | ~ | 2670±601 |
| Mn-ox<sup>g</sup> (mg kg<sup>-1</sup> soil) | 17.8±9.1 | 10.6±4.9 | ~ | 21.9±1.2 |

Group WRB, 2015). <sup>b</sup>Soil texture based on particle size distribution (Soil Science Division Staff, 2017) determined by laser diffraction. <sup>c</sup>Soil organic carbon and total N determined with elemental analyzer. <sup>d</sup>pH (1:10, 24h) determined in 1 mM CaCl<sub>2</sub>.
<sup>e</sup>Effective cation exchange capacity (eCEC) was determined in a 0.0166 M cobalt hexamine (Cohex) extract (ISO, 2007), as the difference between total Co added and Co measured in the extract with inductively coupled plasma-optical emission spectrometry (ICP-OES). <sup>f</sup>Total P determined in aqua-regia digests (Walinga, Lee, Houba, Vark, & Novozamsky, 1995).
<sup>g</sup>Ammonium-oxalate extractive P, Al, Fe and Mn (Schwertmann, 1964).

### Selected rice genotypes

Six upland rice genotypes were selected based on their contrasting performance on low P soils and differences in P acquisition capacity and efficiency (PAE). Detailed information on the selected genotypes is presented in Table 1. Two genotypes with high PAE, DJ 123 and CG 14 (Koide et al., 2013; Mori et al., 2016; Saito et al., 2015), one control genotype with low performance in low P soils, NERICA 4 (De Bauw et al., 2019; Koide et al., 2013; Mori et al., 2016; Pujol & Wissuwa, 2018; Saito et al., 2015) and three other, locally adopted genotypes from Madagascar, Chomrong Dhan, FOFIFA 172 and FOFIFA 173, of which detailed information on their PAE is still lacking (Raboin et al., 2014), were included.

### Field trials

A three-seasonal field experiment (Field experiment 1) was conducted between November and May (2016-2019) at an upland site in Behenjy, located in the central highland region of Madagascar. Details on the locations and standard soil characteristics are presented in Table 2. During the three consecutive years, five (year 1, 2016) or six (years 2-3; 2017-2018) upland rice varieties, two farmyard manure (FYM) treatments (0 and 12.7, 21.7, and 10 t dry weight ha^-1^ respectively for each year) and two mineral P treatments (triple super phosphate (TSP); 0 and 40 kg P ha^-1^) were combined in a full factorial design. The FYM treatment corresponded to an extra addition of respectively 19, 25, 17 kg P ha^-1^ respectively for each year. Four large TSP × FYM treatment blocks (i.e. Ctrl, FYM, TSP, TSP+FYM) were established, in which the varieties were sown (i.e. a nested design). All plots, including control received 80 kg ha^-1^ N as urea and 60 kg ha^-1^ K as K_2_SO_4_. The selected genotypes were sown at a planting distance of 20 cm × 20 cm with two seeds per planting hole. Each genotype was sown in different subplots of 1.92 m^2^ with 12 plants × 4 rows, randomly distributed within the treatment blocks. For the FYM treatments, fresh FYM was locally applied in the planting hole up to ca. 5-10 cm depth and mixed with the soil. For the TSP treatments, TSP granules (2-4 mm diam.) were also locally applied in the planting hole up to ca. 2 cm depth and mixed with the soil. For each year, the FYM was collected from local smallholder farmers; the chemical properties and corresponding nutrient doses of the applied FYM are presented in Table S1 (Supplementary Information). Plots were manually weeded. During the first year, water availability depended on natural rainfall, while in year 2 (2017) and year 3 (2018), the plots were irrigated after each five days without rain, to avoid water stress masking the effects of treatments. Straw and grain yield was determined at maturity (i.e. 138-159 DAS in Year 1 (2016) and 151 DAS in the Year 2 (2017)), by omitting one border row per plot and after sample drying at 70°C. Grain yield is reported at 14% moisture content. Straw and grain samples were ground and composite samples of each genotype were made by combining all subplots within the treatment block for plant analysis. Soil samples were collected right before the start at time zero (i.e. initial, before any treatment) and at 11 (i.e. middle) and 34 months after the first treatment application (i.e. final harvest) in Field 1 (Table 4) as follows. At time zero, the topsoil (0-15 cm depth) was randomly collected from four points covering the field site. The later soil sampling was done after dividing each treatment block into 4 subplots; the same soil depth was applied at the second sampling, and two soil layers (topsoil at 0-15 cm and subsoil at 15-30 cm) were collected at the final sampling. The collected samples from the different points at time zero and from different subplots were separately analysed.

The second field experiment (Field experiment 2) was initiated on a neighbouring field in 2017. This trial was repeated for two seasons, simultaneously with years 2 and 3 of Field experiment 1. All treatments were identical as in year 2 (2017) and year 3 (2018) of the first field experiment. For this second field experiment, a completely randomized design was established by including two replicate blocks of each soil FYM and/or TSP amendment treatment, each having a size of 42.2 m^2^. The six selected rice varieties (Table 1) were planted in subplots which were randomly distributed within each treatment block. Plots were irrigated to field capacity after each five days without rain. All plots received N and K as described for Field experiment 1. Sampling at maturity, straw and grain yield determination, and P analyses occurred similarly as explained for Field experiment 1. Soil samples were collected 21 months after the first treatment application (i.e. final harvest; Table 4) in Field experiment 2, as described for field experiment 1.

A third field experiment (Field experiment 3) was performed to test Lime-TSP-Geno interactions during one season (year 3; 2018). This field trial was established in 2018, having two replicate blocks at the side of Field experiment 2, and included two liming treatments in a completely randomized block design. The combination of six rice genotypes (Table 1) and two liming treatments were randomly assigned over the two blocks. In one treatment 4 t ha^-1^ dolomite was applied (Lime), while in the other treatment 4 t ha^-1^ dolomite and 40 kg P ha^-1^ (as TSP) were amended together (Lime+TSP). All plots received N and K as described for field trial 1. The dolomite was broadcasted and incorporated into the top soil at *ca*. 10-20 cm depth one week before fertilizer application and subsequent sowing. Similar grain and straw data were collected as described for Field experiment 1 and 2. Soil samples were collected 12 months after liming (i.e. final harvest; Table 4) in Field experiment 3, as described for field experiment 1. Soils were collected from two subplots per treatment and analysed separately.

### Soil incubation experiment

A soil incubation experiment was set up to monitor the effects of FYM on soil chemical properties, i.e. pH and 1 mM CaCl_2_ extractable elements. Topsoil (0-20 cm) was collected from the side of Field experiment 1 (Table 2), air-dried and sieved (4 mm). The FYM was collected from the different farmers in 2016, air-dried and ground into 0.2 mm. Characteristics of the FYM used in this experiment are presented in Table S1 (Supplementary Information). The FYM was added at different rates (0, 2.3, 12.0, 23.5, 46.9 g FYM kg^-1^ soil; Table S1) and was homogeneously mixed with the soil. The FYM rates applied on the field ranged 10 to 21.7 ton ha^-1^, and these were more or less equivalent to the intermediate FYM rates used in this incubation experiment (i.e. 12 to 23 g FYM kg^-1^), as the FYM is mixed on the field into the 0-10 cm soil with a bulk density of 0.98 g cm^-3^ (Table S1, Supplementary Information).

Before incubation, the soil and FYM were thoroughly mixed and the soil moisture content was adjusted with distilled water to 20% w/w. The moist soil mixtures (equivalent of 500-g air-dried soil) were placed in a 2-L tray (19.2 cm × 16.4 cm × 9 cm ht.), covered with a plastic lid having pinholes to keep the incubated soil aerated, and placed in a dark incubation room at 25°C. A randomized complete block design was established and three tray replicates were included. These soil-FYM mixtures were not disturbed or mixed during the entire incubation period, to minimize the fluctuation of soil oxygen supply. The soil moisture content was maintained at 20% w/w by spraying distilled water once a week (weight based). Samples were taken from undisturbed soil surface immediately after mixing and after 3, 9, 15, 22, 28, 43, 72, and 99 days.

### Plant and soil analyses

The total P concentrations in the shoots, grains, and FYM were analysed after digestion in concentrated HNO_3_ in a block digester (DigiPREP MS, SCP Science Co.) by inductively coupled plasma optical emission spectrometry (ICP-OES; iCAP 7000 series, Thermo Scientific Inc.). Internal plant reference samples were included in every batch to control quality. Grain and shoot P in each genotype was analysed as a composite sample from each treatment replicate. Total P uptake after harvest was then determined by combining grain and shoot P.

The agronomic efficiency (APE; kg grain kg^-1^ P applied) was calculated as the additional grain yield (kg ha^-^1^^) per unit applied P (kg P ha^-1^) through either or both TSP and/or FYM, when compared to the control plot (Ctrl). For the Lime+P treatment, this was also calculated in comparison with the grain yield of the Lime treatment. None of the control plots (no amendment) had rice grain yields, i.e. the second term was zero in all trials, except for field 3 where rice yield was successful in the limed zero P input.

The soil pH and 1 mM CaCl_2_ extractable elements were measured on equivalent of 3 g air-dried soil, either fresh (incubation) or air-dried (field experiments) that was suspended with 15 ml of 1 mM CaCl_2_ solution. The soil suspensions were equilibrated by end-over-end shaking (26 rpm) for 24h. Next, the samples were centrifuged (2000 rcf, 10 min, 20°C) and the subsample of supernatant was collected. The remaining samples were used for the pH measurement. After additional filtration (0.45 µm), the total elemental concentrations in the collected supernatant, further denoted as CaCl_2_ extractable elements, were measured with inductively coupled plasma mass spectroscopy (ICP-MS; Agilent 7700x, Agilent Technologies, Inc.) after acidification with 1% HNO_3_. The limit of quantification for P was 3 µg L^-1^.

### Statistical analysis

All data were subjected to analysis of variance (ANOVA) using R software, version 3.6.1 (R Development Core Team, 2012). The factors genotype (Geno), FYM and TSP were considered as fixed effects, as also the Liming treatments for Field experiment 3, whereas the subplot replicates were included as a random effect. All field data were analysed separately per field and season, while treatment effects on yield and P uptake were plotted per genotype, including the standard deviation to evaluate the genotypic ranking. The coefficient of variation among genotypes was additionally calculated within each treatment combination.

In order to evaluate the overall effects of genotype, P application and soil acidity and/or FYM, a mixed model fitted on the ensemble of six field trials using JMP software (JMP pro 14, SAS Institute, Inc.), to explain *Y* (i.e. rice grain yield, total P uptake, or APE) as a function of total P applied (*P*_*appl*_, in kg P ha^-1^) and soil acidity, i.e. either CaCl_2_ extractable Al (*Al*, in µg Al L^-1^) or pH as a proxy for the potential mechanism of FYM application. The year of the field trial (i.e. 2016, 2017, 2018) was added as a random variable. The model reads:

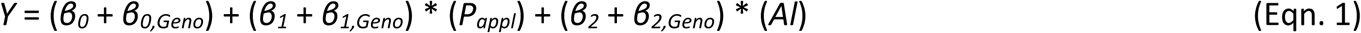

Genotype (*Geno*) is a categorical variable with 6 levels and assigned as the genotype-specific corrector value (*β*_*i,Geno*_, *i is 0, 1 or 2)* to examine the overall effect of genotype and its interaction with P_appl_ and Al. The significance of this genotype-specific correctors denotes whether specific genotypes significantly differ from the mean responses in all six field trials, i.e. which are inherently different (*β*_*0,Geno*_) or different because of its sensitivity to P addition (*β*_*1,Geno*_) or to soil acidity, i.e. Al toxicity (*β*_*2,Geno*_). The incomplete soil sampling did not allow to link soil Al or pH data to corresponding response data and only data of year 3 (i.e. topsoil at final harvest from all three fields; Table 4) were used. Control yields were zero and hence these data were excluded from the modelling, while the data included those of Field 3 (Lime-TSP-Geno) in order to explain the beneficial FYM effects by soil analysis.

## Results

### Genotypic variation in response of rice to FYM and TSP application

Rice grain yield and shoot P uptake largely responded to the application of FYM and TSP in both Field experiments 1 and 2 in all years (Figure 1 & 2). The yield and P uptake were more affected by soil treatments than by genotype. Without P inputs (Ctrl), no grain yield was obtained throughout both field trials (Figure 1), confirming the extremely weathered, P-deficient condition of this soil (Table 2). Rice grain yields reached 6 t ha^-1^ in year 3 (2018) (TSP+FYM) and were lowest in year 1 (2016) when no irrigation was yet adopted. The rice grain yield and total P uptake ranked FYM+TSP>TSP∼FYM>Ctrl, with the FYM either above, similar, or below the TSP, depending on field and year. Interestingly, the average grain yield in the TSP+FYM treatment was consistently higher than the sum of the TSP and FYM treatments separately, suggesting that the effect of the combined application of TSP and FYM was unlikely to be only the mere P addition of each material. Similar trends were observed for P uptake.

**Figure 1:**
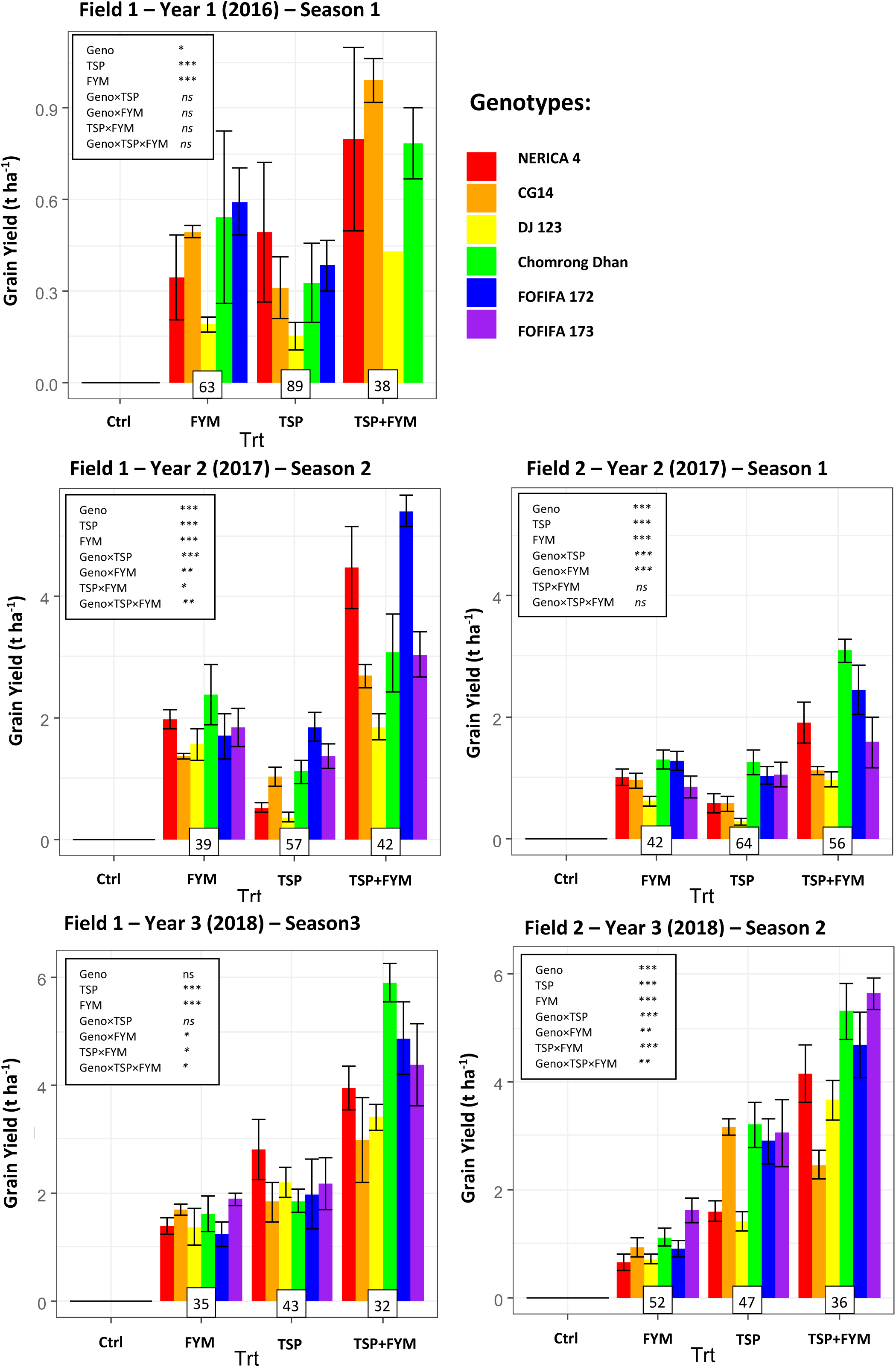
Rice grain yields of Field Experiment 1 and Field Experiment 2, after 1, 2, or 3 seasons of continuous soil fertilization. Barplots are presented as mean with standard deviation. Treatment effects are FYM application (FYM), mineral P fertiliser (TSP), and genotype (Geno), while complete crop failure was observed for the unamended control soil (Ctrl). The coefficient of variation of yield among genotypes is presented for each soil treatment in a box below the bars and was generally largest for the TSP treatment. Note that the scale of the Y-axis is different for each year.

The effects of genotypes on grain yield and P uptake were smaller than soil treatment effects, and were significant (P<0.05) in four of the five trials. The interactions of these endpoints between genotypes and soil treatments were significant in some, but not all trials (Figure 1 & 2). None of the genotypes consistently outperformed others in terms of P uptake or grain yield after a FYM application compared to the mineral P amendment. The ranking of genotypes in terms of total P uptake and grain yield within each treatment was inconsistent over the seasons. The two tropical varieties (i.e. DJ 123 and CG 14) were generally inferior in both grain yield and total P uptake, the control variety NERICA 4 and FOFIFA 173 intermediate, while the two Malagasy highland varieties (i.e. Chomrong Dhan and FOFIFA 172) were superior in all fertilizer treatments (Figures 1 & 2). Interestingly, the coefficient of variation of either grain yield or P uptake among the genotypes within each treatment was highest in the mineral P amended (+TSP) plots compared to those the FYM or the TSP+FYM amended plots in four of the five trials (Figure 1 & 2). This means that FYM amendment decreased the variation in grain yield and P uptake among genotypes. The agronomic P efficiency (APE) of the P applied under the different treatments was consistently highest for the FYM application only (FYM), and lowest for the mineral P amendment (+TSP) (Table 4).

### Genotypic variation in response of rice to lime and TSP application

The limed soil without P input (Lime) had no crop failure and the average grain yield (0.6 t ha^-1^) and total P uptake (0.8 kg P ha^-1^) was similar to that of the FYM treatment in the first season of Field 1 and both seasons of Field 2 (Figure 1, 2, and 3). Combining TSP with lime (Lime+TSP) resulted in a seven-fold increase of grain yield and a six-fold increase of total P uptake, compared to applying lime only (Figure 3). These results are similar to the TSP+FYM treatment from the simultaneously conducted experiment (Field 2, season 2). The liming treatment showed the highest coefficient of variation among genotypes within treatment, clearly due to the good performance of both Chomrong Dhan and FOFIFA 173 in the limed, but strongly P deficient subplots. This coefficient of variation was then strongly reduced upon P fertilization, in combination with lime application (Figure 3). Interestingly, the APE under the Lime+TSP treatment in this experiment was higher than that of the TSP+FYM treatment in experiment 2 (Table 3).

**Table 3.** The agronomic P efficiency (APE) [kg grain yield kg^-1^ P application] of the mineral and organic P amendments in the three field experiments. Mean values with standard error are presented for each treatment, as also the overall mean and coefficient of variation (CV) within treatment.

| Genotype | Field Experiment 1 |  |  |  |  |  |  |  |  |
| --- | --- | --- | --- | --- | --- | --- | --- | --- | --- |
|  | Season 1 (2016) |  |  | Season 2 (2017) |  |  | Season 3 (2018) |  |  |
|  | FYM | TSP | P+FYM | FYM | TSP | P+FYM | FYM | TSP | P+FYM |
| NERICA 4 | 18.2 (7.3) | 12.3 (5.7) | 13.5 (5.1) | 78.8 (6.2) | 13.1 (2.0) | 68.9 (10.3) | 81.8 (8.9) | 70.3 (13.8) | 69.3 (7.2) |
| CG 14 | 26.0 (1.0) | 7.7 (2.5) | 16.8 (1.2) | 55.0 (1.9) | 25.8 (4.0) | 41.4 (2.9) | 99.8 (6.2) | 45.9 (9.1) | 52.5 (13.8) |
| DJ123 | 10.0 (1.3) | 3.8 (1.1) | 7.3 (NA) | 62.4 (10.5) | 9.1 (2.2) | 28.4 (3.2) | 75.1 (23.9) | 54.9 (6.8) | 59.8 (4.1) |
| Chomrong Dhan | 28.4 (15.0) | 8.1 (3.2) | 13.3 (1.9) | 95.4 (19.7) | 27.7 (4.6) | 47.2 (10.0) | 95.2 (19.3) | 46.5 (5.5) | 103.7 (6.3) |
| FOFIFA 172 | 31.3 (6.0) | 9.6 (2.1) | NA | 67.7 (15.0) | 46.2 (5.8) | 83.1 (4.0) | 72.6 (13.9) | 49.5 (16.3) | 85.6 (12.0) |
| FOFIFA 173 | NA | NA | NA | 73.8 (12.4) | 34.4 (5.2) | 46.7 (5.7) | 110.7 (7.1) | 54.2 (12.1) | 77.0 (13.5) |
| Overall Mean (SE) | 23.0 (3.4) | 8.5 (1.8) | 14.2 (1.7) | 72.5 (5.3) | 26.4 (2.9) | 54.6 (4.4) | 88.0 (6.3) | 53.2 (4.4) | 76.2 (4.7) |
| CV | 63.3 | 88.5 | 37.7 | 38.6 | 57.4 | 41.9 | 38.1 | 43.0 | 32.4 |

| Genotype | Field Experiment 2 |  |  |  |  |  | Field Experiment 3 |  |
| --- | --- | --- | --- | --- | --- | --- | --- | --- |
|  | Season 1 (2017) |  |  | Season 2 (2018) |  |  | Season 1 (2018) |  |
|  | FYM | TSP | P+FYM | FYM | TSP | P+FYM | Lime+TSP vs ctrl | Lime+TSP vs Liming |
| NERICA 4 | 40.6 (5.4) | 14.3 (4.0) | 29.1 (5.2) | 38.8 (9.3) | 40.1 (4.5) | 72.9 (9.2) | 87.9 (12.6) | NA |
| CG 14 | 38.4 (4.9) | 14.2 (2.9) | 17.2 (1.0) | 55.0 (10.5) | 79.1 (3.8) | 43.3 (4.7) | 133.2 (41.3) | 123.2 (41.3) |
| DJ123 | 24.5 (3.0) | 7.1 (1.4) | 14.9 (1.8) | 41.8 (5.5) | 35.2 (4.3) | 64.3 (6.5) | 60.6 (13.6) | 48.9 (13.6) |
| Chomrong Dhan | 51.9 (6.2) | 31.2 (5.1) | 47.6 (2.9) | 65.7 (9.3) | 80.1 (10.7) | 93.3 (9.1) | 123.0 (15.9) | 81.3 (15.9) |
| FOFIFA 172 | 50.8 (6.3) | 25.9 (3.8) | 37.7 (6.2) | 53.4 (8.6) | 72.5 (10.4) | 82.1 (10.7) | 176.0 (13.5) | 168.5 (13.5) |
| FOFIFA 173 | 34.2 (7.1) | 26.4 (5.1) | 24.4 (6.4) | 95.7 (13.4) | 76.3 (15.3) | 99.0 (5.1) | 111.9 (29.2) | 92.3 (29.2) |
| Overall Mean (SE) | 41.0 (2.6) | 20.4 (2.0) | 28.9 (2.4) | 57.2 (4.5) | 62.2 (4.5) | 75.5 (4.2) | 109.4 (10.2) | 94.4 (13.1) |
| CV | 42.0 | 64.3 | 55.7 | 52.1 | 47.3 | 36.1 | 40.4 | 52.1 |

**Figure 2:**
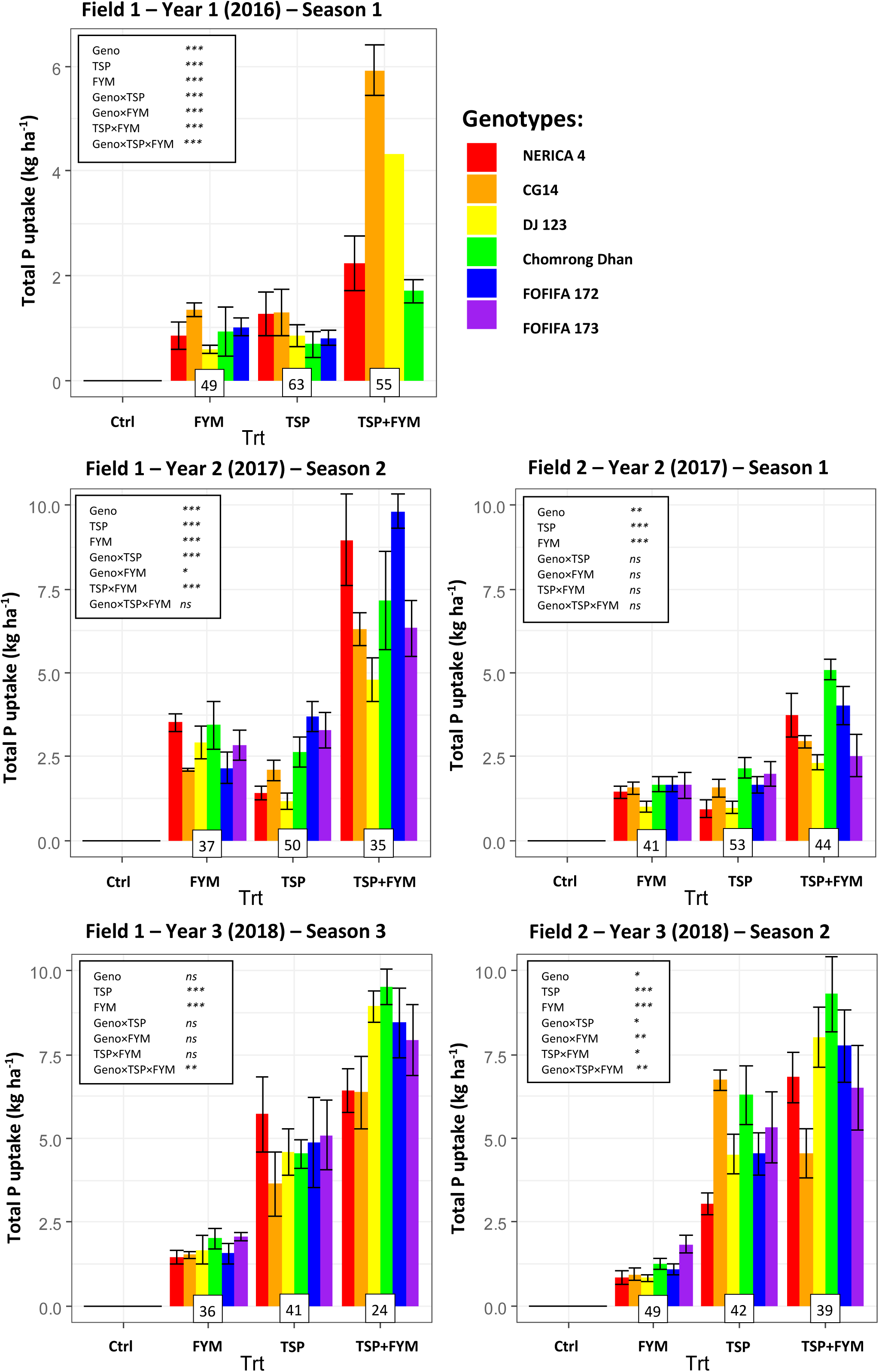
The total P uptake in rice grain and straw at maturity in Field Experiment 1 and Field Experiment 2, after 1, 2, and 3 years of continuous soil fertilization. Barplots are presented as mean with standard deviation. Treatment effects are FYM application (FYM), mineral P amendment (TSP), and genotype selection (Geno), while complete crop failure was observed for the unamended control soil (Ctrl). The coefficient of variation of the total P uptake among genotypes is presented for each soil treatment in a box below the bars and was generally largest for the TSP treatment P. Note that the scale of the Y-axis is different for each year.

**Figure 3:**
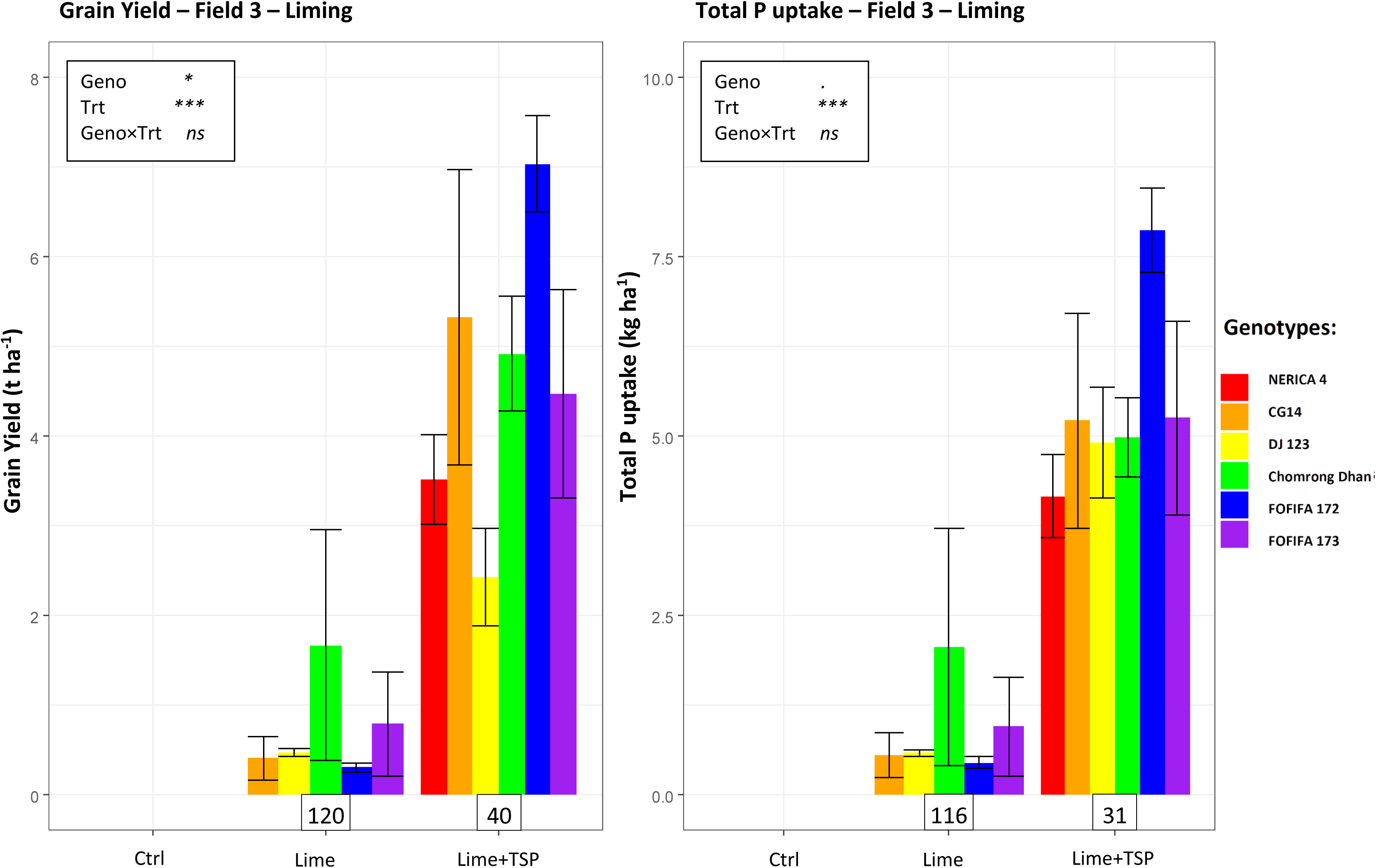
The grain yield and total P uptake at maturity in Field Experiment 3 (Lime and Lime+TSP). Barplots are presented as mean with standard deviation. The coefficient of variation among genotypes is given for each treatment in a box below the bars.

### Changes in soil properties to FYM and lime applications

The FYM applications to the soil in the laboratory incubation increased the soil pH from an initial value of 4.3 up to 4.5-6.1, depending on the application rate. The sharp and immediate increase in pH was followed by a gradual pH decrease but all soils amended with FYM had higher pH than the unamended soil (Figure 4a). The FYM applications increased the CaCl_2_-extractable P (Figure 4b) and decreased CaCl_2_-extractable Al up to pH 5.0 (Figure 4c). The CaCl_2_-extractable Al exhibited a distinct amphoteric behaviour to soil pH and to FYM dose, i.e. soluble Al decreased at initial FYM doses and increased again at the two highest FYM rates for which pH increased to ca. 5.5 and 6.0 (Figure 4c). The CaCl_2_ extractable P increased with increasing FYM doses and the semi-log trend to pH was linear (Figure 4b).

**Figure 4:**
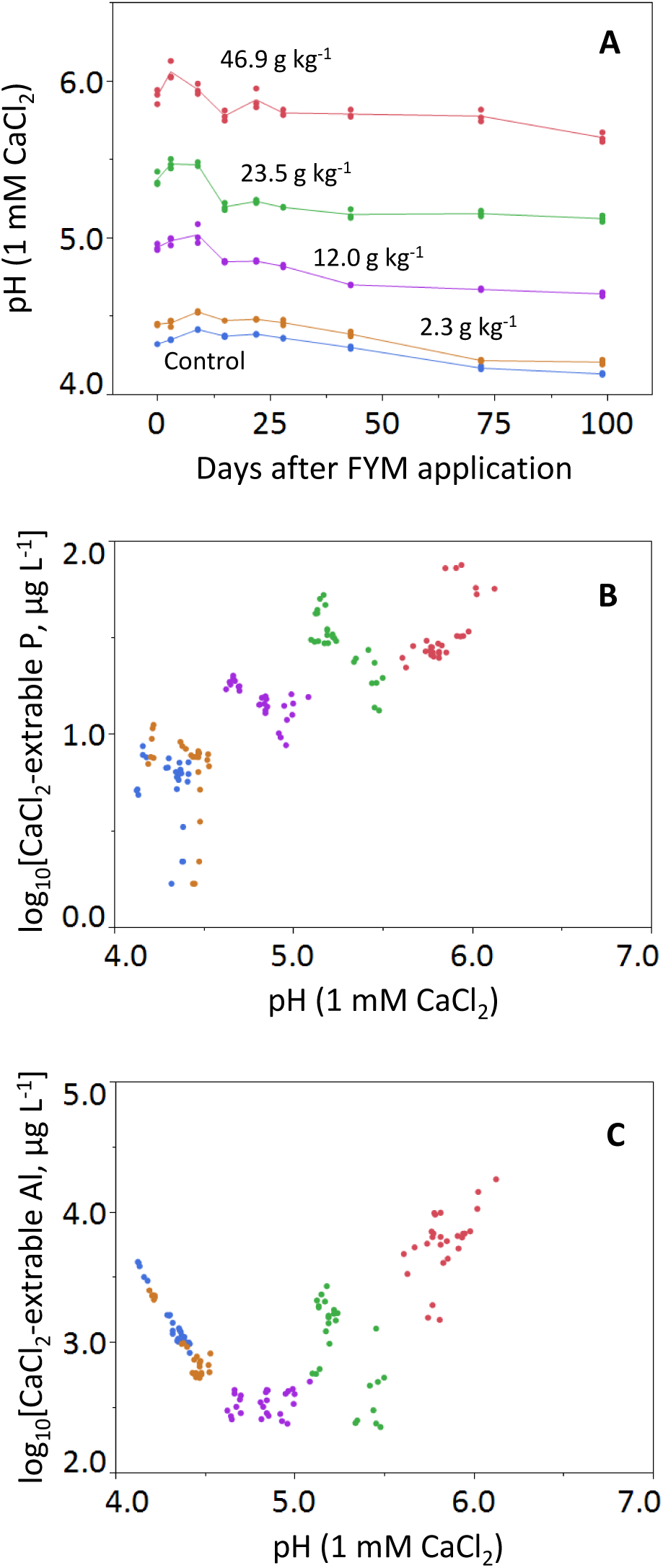
The dynamics of (a) the soil pH (top), (b) 1 mM CaCl_2_ extractable P (middle), and (c) 1 mM CaCl_2_ extractable Al (bottom) of the soils incubated in laboratory conditions. Note the log scale of extractable solute concentrations. Soils were amended with four rates of FYM (0, 2.3, 12.0, 23.5, 46.9 g FYM kg^-1^ soil, n=3 each), and incubated at 25 °C without leaching.

Similar, but less pronounced effects were observed in soils of the field trials. In the control treatments, the soil pH at final harvest decreased over time and CaCl_2_ extractable Al increased (Table S2, Supplementary Information). In Field 1, initially, both FYM and TSP treatments increased pH and decreased extractable Al, while after the final harvest, only such changes were found in the FYM treatments (FYM and TSP+FYM). These changes were observed in both surface and subsoils. Similar trends were observed in Field 2 except that the pH increase and extractable Al decrease was consistently similar in all treatments. The CaCl_2_-extractable P showed a clear increase compared to the control after either TSP and/or FYM application; however, no additive effect was observed in the TSP+FYM treatment compared to the single TSP and FYM treatment (Table S2, Supplementary Information). Liming strongly increased soil pH and the CaCl_2_-extractable P, and it strongly reduced the CaCl_2_-extractable Al, even in the subsoil (Table S2, Supplementary Information). Both liming only as the combination of lime with TSP (i.e. Lime+TSP) resulted in the highest CaCl_2_-extractable P concentrations of all treatments, taking to account that samples were taken only 12 months after application. The CaCl_2_ extractable P in these soils was never above 25 µg P L^-1^ whereas 100 µg P L^-1^ is considered as a threshold for adequate P supply (De Bauw et al., 2019; Six et al., 2014).

### Overall genotypic effects to P addition and soil acidity

Equation (1) was fitted to the ensemble of six field trials and revealed the contribution of Geno, TSP and soil acidity to the overall crop responses (Table S 3 and Table 4). The effects of including either soil pH or CaCl_2_ extractable Al were similar to explain grain yield, P uptake or APE. Only the Al based data, rather than soil pH, were further used to indicate effects of Al toxicity. The overall variation in rice grain yield explained by soil and genotypic factors rank P addition>> genotype> soil acidity>genotype interaction with P addition or soil acidity (Table 4). By including the zero yielding control, the effects of genotype became even smaller in the mixed models (details not shown).

**Table 4.** Summary of rice grain yield, shoot P uptake, and agronomic P efficiency (APE) in the six field trials. Data include all treatments (including lime treatments) but exclude the zero yielding control treatments. Data show statistical comparison (Tuckey’s HSD) for the least square means of genotypes after fitting model Eqn. (1) that is presented in Table S2 (Supplementary Information). The p-values are assigned as ***, p<0.001; **, p<0.01; *, p<0.05. Means in the same column followed by the same character are not significantly different (Tukey’s HSD, p<0.05). The effect size (% variance explained) was calculated by including year as fixed (not random) factor, while the models presented in Eqn(1) (Table S2) present a mixed modelling with year as random factor.

| Genotype | Grain yield<br>[t ha<sup>-1</sup>] | P uptake<br>[kg P ha<sup>-1</sup>] | Agronomic P efficiency (APE)<br>[kg grain yield<br>kg<sup>-1</sup> P application] |
| --- | --- | --- | --- |
| DJ 123 | 1.1<sup>d</sup> | 2.8<sup>b</sup> | 28<sup>b</sup> |
| CG 14 | 1.3<sup>cd</sup> | 2.8<sup>b</sup> | 38<sup>b</sup> |
| NERICA 4 | 1.6<sup>bc</sup> | 2.8<sup>b</sup> | 39<sup>b</sup> |
| FOFIFA 173 | 1.9<sup>ab</sup> | 3.2<sup>ab</sup> | 52<sup>a</sup> |
| FOFIFA 172 | 2.1<sup>a</sup> | 3.5<sup>ab</sup> | 51<sup>a</sup> |
| Chomrong Dhan | 2.2<sup>a</sup> | 3.7<sup>a</sup> | 55<sup>a</sup> |
| Factor | % variance explained |  |  |
| Year | 18.0*** | 10.9*** | 31.9*** |
| P application (kg ha<sup>-1</sup>) | 28.5*** | 43.2*** | 0.1<sup>n.s.</sup> |
| CaCl<sub>2</sub>-extractable Al in soil (µg L<sup>-1</sup>) | 1.4*** | <0.1<sup>n.s.</sup> | 2.2*** |
| Geno | 5.6*** | 1.4* | 7.3*** |
| Geno × P<sub>appl</sub> | 2.6*** | 1.5* | 1.7* |
| Geno × Al | 1.0* | 0.5<sup>n.s.</sup> | 1.2<sup>n.s.</sup> |
| Total model R<sup>2</sup> | 0.57 | 0.57 | 0.45 |

The model was extended to test if the FYM dose also affected crop response beyond its effect on reducing soil acidity. This was done by extending Eqn. (1) with the added FYM dose and its interaction with Geno for all field data of year 3, i.e. the TSP-FYM data of Fields 1 & 2 and the Lime trial of Field 3. That showed that FYM had no statistically significant effect on grain yield after correction for effects of its P input and reducing soil acidity. This result suggests that the main effect of FYM here is through its effect on soil acidity and Al.

In the ensemble of these data, genotypes Chomrong Dhan and FOFIFA 172 performed significantly superior whereas DJ 123 and CG 14 were significantly lower (Table S3 and Table 4). The coefficients for the genotype-specific corrector values for either P addition rate or CaCl_2_ extractable Al suggest that FOFIFA 172 is more responsive to P addition than Chomrong Dhan and that the latter is more sensitive to Al (Table S3). That is also clearly visualized in the liming trial where Chomrong Dhan performs well under low P if the soil acidity constraint is lifted (Figure 3 versus Figure 2).

## Discussion

### Yield and P uptake of upland rice after FYM application

This study could not identify genotypes that consistently outperform others after the application of organic matter, in contrast to only mineral P amendments, or vice versa. The main significant effects of genotypes (Table 4) are more affected by their growth potential, i.e. under TSP+FYM application, rather than by their inherent differences in in PAE and PUE that exist among rice genotypes (De Bauw et al., 2019; Nestler and Wissuwa, 2016; Vandamme et al., 2016a). This indicates that genotype selection likely becomes inferior to management options when soil limitations are very severe. It is interesting that the coefficient of variation of yield and P uptake generally decreases after FYM application compared to mineral P amendment only (Figures 1 & 2); and that it further decreases for yield when combining FYM with a mineral P amendment. This can be explained by the fact that the genetic variability under a mineral P amendment includes sensibility of rice genotypes to multiple factors (e.g. low pH, P deficiency, cold tolerance), while FYM application ameliorates Al toxicity (Haynes & Mokolobate, 2001). However, when only FYM is applied, the genotypic variation still includes tolerance to P deficiency. Hence, when both FYM and mineral P are combined, yields are closer to their potential, with fewer environmental limitations contributing to the genotypic variation (i.e. cold tolerance) (Raboin et al., 2014).

Under combined TSP and FYM amendments, the genotypes Chomrong Dhan and FOFIFA 172 generally outperformed in terms of yields, which can be explained by their improved tolerance to cold conditions; and possibly explaining why these genotypes are widely adopted by Malagasy farmers (Raboin et al., 2014). Additionally, it was suggested that Chomrong Dhan has a high PAE, as it gives high grain yields under low input system (Raboin et al., 2014), but such a high P acquisition under P limitations could not consistently be confirmed for Chomrong Dhan. The multivariate model Eqn (1) fitted to the ensemble of data allows to interpret the genotypic variation and its interaction with treatments, i.e. quantify the so-called G × E interactions (Table S3). This showed that the most performant genotype (Chomrong Dhan) is most sensitive to soil acidity whereas the other performant FOFIFA 172 is relative more tolerant; conversely, the sensitivity to P input shows that responsiveness to P input is in the reverse order, i.e. FOFIFA 172 is more responsive (less performant under low P input) than Chomrong Dhan. The generally P-efficient genotype DJ123, known for its good performance on low P soils (Mori et al., 2016; Nestler & Wissuwa, 2016; Vandamme, Wissuwa, et al., 2016), was not high yielding in the low P environment of this study and it even showed comparable or lower yields than the generally P-inefficient rice genotype, NERICA 4. Hence, this indicates that this P-efficient genotype (DJ123) is likely not adapted to the cold highlands of Madagascar while NERICA 4 seems somehow more robust in such environment. It was previously observed that DJ123 was extremely sensitive to Al toxicity (Famoso et al., 2011), while NERICA 4 was more moderately tolerant (Kang et al., 2012); and no such differences in Al sensitivity among these genotypes were found here as judged from the statistical analysis (Table S3, Supplementary Information).

### Effects of FYM application on P and Al dynamics in weathered soils and implications for rice yields and P uptake

This study demonstrates that FYM application improve yields up to above 5 t ha^-1^ when additionally amended with mineral P fertilizer, and positive interactions between FYM and TSP were found. The benefits of FYM applications confirmed earlier data for rice (Andriamananjara et al., 2018; Haefele et al., 2006; Satyanarayana et al., 2002), however, the mechanisms were never confirmed. Andriamananjara *et al*. (2018) suggested that the benefits of FYM application to weathered upland soils relate to the better water supply and reducing Al toxicity. By including irrigation here in Years 2 & 3 of the study, the effect of FYM on water availability was excluded. This study now confirms that the application of FYM initially increases soil pH, which concomitantly reduces soluble Al and increases P availability. An initial strong liming effect of FYM application is observed in the soil incubation trial (Figure 4) but, interestingly, on the field these liming effects of FYM application on soil pH are much smaller on the longer term. This suggests that the initial pH increase over the first two or three months of residue decomposition (Haynes & Mokolobate, 2001) is followed by a fast pH decrease because of mineralisation processes, i.e. nitrification, and subsequent leaching of nitrates and/or neutral cations on the field (Rowell, 2014). These observations confirm that pH effects after FYM application are indeed local and short term (Cong & Merckx, 2005), leaving a short-term window of opportunity for crop production. In contrast to the short term pH effects, the reduced Al availability and increased P availability after FYM application are more sensitive indices of a longer term liming effect because small pH changes in the range 4-5 affect soluble Al by over one order of magnitude (Figure 4b). The amphoteric behaviour of Al solubility likely indicates the trend of reduced Al^3+^ in the most acid part followed by solubilisation of Al complexed with dissolved organic matter at high FYM doses and higher pH. The increased P availability on the longer term after FYM application likely follows from (i) ligand exchange (i.e. organic sorbates competing with P for mineral binding sites), and, (ii) the slow release of organic P by mineralization. (Guppy et al., 2005; Pypers et al., 2005)

Here, we now argue that the positive FYM-TSP interaction effects in acid upland soils are related to mitigating Al toxicity. This is supported by the inclusion of a liming field trial and the statistical observation that soluble Al and P dose explain crop response, i.e. that the FYM dose does not further explain the effects across trials with lime and FYM. The local and short term pH increase after FYM application and the associated reduction in Al toxicity allows a strong initial root proliferation during seedling establishment, thereby enhancing an increased P uptake from the TSP granules (Andriamananjara et al., 2018; Cong & Merckx, 2005), so further enhancing root growth, P uptake, plant establishment, and final yield. A hydroponic study reported a critical Al concentration of 1350 μg Al L^-1^ for rice (Zhang et al., 2016). In our field experiments, the measured CaCl_2_-extractable Al generally exceeded this limit, although the concentration tended to be lower in the FYM and liming treatments (Table S2, Supporting Information). In the year 3, these concentrations were reduced by FYM application as well as liming treatment.

### The agronomic consequences of FYM applications and implications to field management

Rice genotype selection is thus inferior to farm management effects on weathered acid soils when considering yields, but depending on the environment, selecting cold-, drought-, and/or disease tolerant varieties can help to further improve the yield potential after tackling soil limitations by additions of FYM and TSP. It is striking that attempting rice cultivation on such acid soils without FYM or mineral P amendments (i.e. control treatment) even further decreases soil pH and increases Al in soil solution compared to the initial condition (Table 4). This is explained by (i) the natural tendency for soil acidification over time, (ii) an increased aeration and enhanced microbial activity due to soil cultivation, and (iii) the nitrification of the applied urea (Rowell, 2014). Farmers should thus recognize that cultivation and application of urea on such already acid soils, without other amendments further aggravates Al toxicity to the rice plant. This work clearly demonstrates how liming facilitates rice cultivation on highly weathered soils, but without additional nutrient inputs yields remain very low (Figure 3). Liming only is thus not a sustainable way to increase upland rice production, while liming combined with mineral P amendments quickly and strongly increase rice yields. It should however be regarded that both lime and TSP are often beyond reach of farmers, following market availability and costs (AGRA, 2019; Chianu et al., 2012; Raboin et al., 2016).

As FYM is often more easily accessible to farmers, the application of FYM forms a huge opportunity to more sustainably increase rice yields on weathered acid low P soils. However, in order to obtain short-term soil pH effects on the field, this work indicates that at least rates of 10 t ha^-1^ of FYM are required (Figure 4), or farmers should strongly concentrate the FYM in the planting hole to facilitate local pH effects close to the seedling. Applying FYM to uncultivatable fields thus enables rice production, but rice yields still remain very low (below 2 t ha^-1^ even after three years of FYM application), following remaining P limitations as the applied P in the FYM is not readily available to the plant. Therefore, it was stated before and confirmed here that the use of organic amendments alone are unlikely to overcome P deficiency in such strongly depleted soils (Andriamananjara et al., 2018; Nziguheba, 2007; Nziguheba et al., 2016). Hence, the combination of organic resources with mineral fertilizers would offer a more sustainable way for increasing soil fertility and rice production, while reducing the use of inaccessible mineral fertilizer. (Buresh et al., 1997; Chivenge et al., 2009; Nziguheba et al., 2016; Six et al., 2014).

## Acknowledgements

This work was financed by a C1 project (C16/15/042), funded by the KU Leuven. We thank Maarten Everaert, Mino Nandrianina Rakotonandrasana, Tsanta Raharijaona, and Seheno Rinasoa for their assistance during the field experiments. We further thank Marie-Paule Razafimanantsoa and all other LRI and KU Leuven staff who assisted in this work. We are grateful to Karlien Cassaert for the assistance in administration.

## Notes

### Competing Interest Statement

The authors have declared no competing interest.

## References

AGRA. (2019). Feeding Africa’s soils: Fertilizers to support Africa’s agricultural transformation.

Nairobi, Kenya. Andriamananjara, A., Rakotoson, T., Razanakoto, O. R., Razafimanantsoa, M.-P., Rabeharisoa, L., & Smolders, E. (2018). Farmyard manure application in weathered upland soils of Madagascar sharply increase phosphate fertilizer use efficiency for upland rice. Field Crops Research, 222, 94–100. doi: 10.1016/j.fcr.2018.03.022

Andriamananjara, Andry, Rakotoson, T., Razafimbelo, T., Rabeharisoa, L., Razafimanantsoa, M.-P., & Masse, D. (2019). Farmyard manure improves phosphorus use efficiency in weathered P deficient soil. Nutrient Cycling in Agroecosystems. doi: 10.1007/s10705-019-10022-3

Buresh, R. J., Sanchez, P. A., Calhoun, F., Palm, C. A., Myers, R. J. K., & Nandwa, S. M. (1997). Combined Use of Organic and Inorganic Nutrient Sources for Soil Fertility Maintenance and Replenishment. doi: 10.2136/sssaspecpub51.c8

Chauhan, B. S., Jabran, K., & Mahajan, G. (Eds.). (2017). Rice Production Worldwide. doi: 10.1007/978-3-319-47516-5

Chianu, J. N., Chianu, J. N., & Mairura, F. (2012, April). Mineral fertilizers in the farming systems of sub-Saharan Africa. A review. Agronomy for Sustainable Development, Vol. 32, pp. 545–566. doi: 10.1007/s13593-011-0050-0

Chivenge, P., Vanlauwe, B., Gentile, R., Wangechi, H., Mugendi, D., van Kessel, C., & Six, J. (2009). Organic and mineral input management to enhance crop productivity in central Kenya. Agronomy Journal, 101(5), 1266–1275. doi: 10.2134/agronj2008.0188x

Clark, R. B. (1983). Plant genotype differences in the uptake, translocation, accumulation, and use of mineral elements required for plant growth. Plant and Soil, Vol. 72, pp. 175–196. doi: 10.2307/42935183

Cong, P. T., & Merckx, R. (2005). Improving phosphorus availability in two upland soils of Vietnam using shape Tithonia diversifolia H. Plant and Soil, 269(1–2), 11–23. doi: 10.1007/s11104-004-1791-1

De Bauw, P., Vandamme, E., Lupembe, A., Mwakasege, L., Senthilkumar, K., & Merckx, R. (2019). Architectural root responses of rice to reduced water availability can overcome phosphorus stress. Agronomy, 9(1). doi: 10.3390/agronomy9010011

Diagne, A., Alia, D. Y., Amovin-Assagba, E., Wopereis, M. C. S., Saito, K., & Nakelse, T. (2013). Farmer perceptions of the biophysical constraints to rice production in sub-Saharan Africa, and potential impact of research. In Realizing Africa’s rice promise (pp. 46–68). doi: 10.1079/9781845938123.0046

Famoso, A. N., Zhao, K., Clark, R. T., Tung, C. W., Wright, M. H., Bustamante, C., … McCouch, S. R. (2011). Genetic architecture of aluminum tolerance in rice (oryza sativa) determined through genome-wide association analysis and qtl mapping. PLoS Genetics, 7(8). doi: 10.1371/journal.pgen.1002221

Guppy, C. N., Menzies, N. W., Moody, P. W., & Blamey, F. P. C. (2005, January 1). Competitive sorption reactions between phosphorus and organic matter in soil: A review. Australian Journal of Soil Research, Vol. 43, pp. 189–202. doi: 10.1071/SR04049

Haefele, S. M., Naklang, K., Harnpichitvitaya, D., Jearakongman, S., Skulkhu, E., Romyen, P., … Wade, L. J. (2006). Factors affecting rice yield and fertilizer response in rainfed lowlands of northeast Thailand. Field Crops Research, 98(1), 39–51. doi: 10.1016/j.fcr.2005.12.003

Haynes, R. J., & Mokolobate, M. S. (2001). Amelioration of Al toxicity and P deficiency in acid soils by additions of organic residues: a critical review of the phenomenon and the mechanisms involved. In Nutrient Cycling in Agroecosystems (Vol. 59).

ISO. (2007). ISO 23470:Soil quality — Determination of effective cation exchange capacity (CEC) and exchangeable cations using a hexamminecobalt trichloride solution. Retrieved from https://www.iso.org/obp/ui/#iso:std:iso:23470:ed-1:v1:en

IUSS Working Group WRB. (2015). World reference base for soil resources, update 2015 International soil classification system for naming soils and creating legends for soil maps. Rome.

Kang, D.-J., Futakuchi, K., Seo, Y.-J., Vijarnsorn, P., & Ishii, R. (2012). Evaluation of Al-tolerance on upland and lowland types of NERICA lines under hydroponic conditions. Journal of Crop Science and Biotechnology, 15(1), 25–31. doi: 10.1007/s12892-011-0083-6

Koide, Y., Pariasca Tanaka, J., Rose, T., Fukuo, A., Konisho, K., Yanagihara, S., … Wissuwa, M. (2013). Qtls for phosphorus deficiency tolerance detected in upland nerica varieties. Plant Breeding, 132(3), 259–265. doi: 10.1111/pbr.12052

Mehra, P., Pandey, B. K., & Giri, J. (2017). Improvement in phosphate acquisition and utilization by a secretory purple acid phosphatase (OsPAP21b) in rice. Plant Biotechnology Journal, 15(8), 1054–1067. doi: 10.1111/pbi.12699

Mori, A., Fukuda, T., Vejchasarn, P., Nestler, J., Pariasca-Tanaka, J., & Wissuwa, M. (2016). The role of root size versus root efficiency in phosphorus acquisition in rice. Journal of Experimental Botany, 67(4), 1179–1189. doi: 10.1093/jxb/erv557

Mueller, N. D., Gerber, J. S., Johnston, M., Ray, D. K., Ramankutty, N., & Foley, J. A. (2012). Closing yield gaps through nutrient and water management. Nature, 490(7419), 254–257. doi: 10.1038/nature11420

Nestler, J., & Wissuwa, M. (2016). Superior Root Hair Formation Confers Root Efficiency in Some, But Not All, Rice Genotypes upon P Deficiency. Frontiers in Plant Science, 7, 1935. doi: 10.3389/fpls.2016.01935

Nishigaki, T., Tsujimoto, Y., Rinasoa, S., Rakotoson, T., Andriamananjara, A., & Razafimbelo, T. (2019). Phosphorus uptake of rice plants is affected by phosphorus forms and physicochemical properties of tropical weathered soils. Plant and Soil, 435(1–2), 27–38. doi: 10.1007/s11104-018-3869-1

Nziguheba, G. (2007). Overcoming phosphorus deficiency in soils of Eastern Africa: recent advances and challenges. In Advances in Integrated Soil Fertility Management in sub-Saharan Africa: Challenges and Opportunities (pp. 149–160). doi: 10.1007/978-1-4020-5760-1_13

Nziguheba, G., Zingore, S., Kihara, J., Merckx, R., Njoroge, S., Otinga, A., … Vanlauwe, B. (2016). Phosphorus in smallholder farming systems of sub-Saharan Africa: implications for agricultural intensification. Nutrient Cycling in Agroecosystems, 104(3), 321–340. doi: 10.1007/s10705-015-9729-y

Osborne, L. D., & Rengel, Z. (2002). Growth and P uptake by wheat genotypes supplied with phytate as the only P source. Australian Journal of Agricultural Research, 53(7), 845–850. doi: 10.1071/AR01102

Pujol, V., & Wissuwa, M. (2018). Contrasting development of lysigenous aerenchyma in two rice genotypes under phosphorus deficiency. BMC Research Notes, 11(1), 60. doi: 10.1186/s13104-018-3179-y

Pypers, P., Verstraete, S., Thi, C. P., & Merckx, R. (2005). Changes in mineral nitrogen, phosphorus availability and salt-extractable aluminium following the application of green manure residues in two weathered soils of South Vietnam. Soil Biology and Biochemistry, 37(1), 163–172. doi: 10.1016/j.soilbio.2004.06.018

R Development Core Team. (2012). R: A Language and Environment for Statistical Computing. Vienna, Austria: R Foundation for Statistical Computing.

Raboin, L.-M., Randriambololona, T., Radanielina, T., Ramanantsoanirina, A., Ahmadi, N., & Dusserre, J. (2014). Upland rice varieties for smallholder farming in the cold conditions in Madagascar’s tropical highlands. Field Crops Research, 169, 11–20. doi: 10.1016/J.FCR.2014.09.006

Raboin, L. M., Razafimahafaly, A. H. D., Rabenjarisoa, M. B., Rabary, B., Dusserre, J., & Becquer, T. (2016). Improving the fertility of tropical acid soils: Liming versus biochar application? A long term comparison in the highlands of Madagascar. Field Crops Research, 199, 99–108. doi: 10.1016/j.fcr.2016.09.005

Rakotoson, T. (2014). Overcoming phosphate deficiency in flooded rice in Madagascar.

Richardson, A. E., Lynch, J. P., Ryan, P. R., Delhaize, E., Smith, F. A., Smith, S. E., … Simpson, R. J. (2011). Plant and microbial strategies to improve the phosphorus efficiency of agriculture. Plant and Soil, 349(1–2), 121–156. doi: 10.1007/s11104-011-0950-4

Rose, T. J., Impa, S. M., Rose, M. T., Pariasca-Tanaka, J., Mori, A., Heuer, S., … Wissuwa, M. (2013). Enhancing phosphorus and zinc acquisition efficiency in rice: a critical review of root traits and their potential utility in rice breeding. Annals of Botany, 112(2), 331–345. doi: 10.1093/aob/mcs217

Rowell, D. L. (2014). Soil Science : Methods & Applications. Routledge.

Saito, K., Vandamme, E., Segda, Z., Fofana, M., & Ahouanton, K. (2015). A Screening Protocol for Vegetative-stage Tolerance to Phosphorus Deficiency in Upland Rice. Crop Science, 55(3), 1223. doi: 10.2135/cropsci2014.07.0521

Satyanarayana, V., Vara Prasad, P. V., Murthy, V. R. K., & Boote, K. J. (2002). Influence of integrated use of farmyard manure and inorganic fertilizers on yield and yield components of irrigated lowland rice. Journal of Plant Nutrition, 25(10), 2081–2090. doi: 10.1081/PLN-120014062

Schwertmann, U. (1964). Differenzierung der Eisenoxide des Bodens durch Extraktion mit Ammoniumoxalat-Lösung. Zeitschrift Für Pflanzenernährung, Düngung, Bodenkunde, 105(3), 194–202. doi: 10.1002/jpln.3591050303

Six, L., Smolders, E., & Merckx, R. (2014). Testing phosphorus availability for maize with DGT in weathered soils amended with organic materials. Plant and Soil, 376(1), 177–192. doi: 10.1007/s11104-013-1947-y

Soil Science Division Staff. (2017). Soil Survey Manual (USDA Handb; C. Ditzler, K. Scheffe, & H. C. Monger, Eds.). Washington D.C.: Government Printing Office.

Stoorvogel, J. J., Smaling, E. M. A., & Janssen, B. H. (1993). Calculating soil nutrient balances in Africa at different scales. Fertilizer Research, 35(3), 227–235. doi: 10.1007/BF00750641

Thibaud, M. C., Morel, C., & Fardeau, J. C. (1988). Contribution of phosphorus issued from crop residues to plant nutrition. Soil Science and Plant Nutrition, 34(4), 481–491. doi: 10.1080/00380768.1988.10416464

Turner, B. L. (2006). Organic phosphorus in Madagascan rice soils. Geoderma, 136(1–2), 279–288. doi: 10.1016/j.geoderma.2006.03.043

Vandamme, E., Rose, T., Saito, K., Jeong, K., & Wissuwa, M. (2016). Integration of P acquisition efficiency, P utilization efficiency and low grain P concentrations into P-efficient rice genotypes for specific target environments. Nutrient Cycling in Agroecosystems, 104(3), 413–427. doi: 10.1007/s10705-015-9716-3

Vandamme, E., Wissuwa, M., Rose, T., Dieng, I., Drame, K. N., Fofana, M., … Saito, K. (2016). Genotypic Variation in Grain P Loading across Diverse Rice Growing Environments and Implications for Field P Balances. Frontiers in Plant Science, 7, 1435. doi: 10.3389/fpls.2016.01435

Walinga, I., Lee, J. J., Houba, V. J. G., Vark, W., & Novozamsky, I. (1995). Plant Analysis Manual. Springer Netherlands.

Wissuwa, M., & Ae, N. (2001). Genotypic variation for tolerance to phosphorus deficiency in rice and the potential for its exploitation in rice improvement. Plant Breeding, 120(1), 43–48. doi: 10.1046/j.1439-0523.2001.00561.x

Zhang, P., Zhong, K., Tong, H., Shahid, M. Q., & Li, J. (2016). Association mapping for aluminum tolerance in a core collection of rice landraces. Frontiers in Plant Science, 7(OCTOBER2016). doi: 10.3389/fpls.2016.01415

